# Quantifying uncertainty in Protein Representations Across Models and Task

**DOI:** 10.1101/2025.04.30.651545

**Authors:** R Prabakaran, Y Bromberg

**Affiliations:** Department of Biology, Emory University, Atlanta, GA 30322, USA; Department of Computer Science, Emory University, Atlanta, GA 30322, USA

**Keywords:** Language models, Artificial Intelligence, Embeddings, pLMs, Prediction confidence, Evaluation

## Abstract

Embeddings, derived by language models, are widely used as numeric proxies for human language sentences and structured data. In the realm of biomolecules, embeddings serve as efficient sequence and/or structure representations, enabling similarity searches, structure and function prediction, and estimation of biophysical and biological properties. However, relying on embeddings without assessing the model’s confidence in its ability to accurately represent molecular properties is a critical flaw—akin to using a scalpel in surgery without verifying its sharpness.

In this study, we propose a means to evaluate the ability of protein language models to represent proteins, assessing their capacity to encode biologically relevant information. Our findings reveal that low-quality embeddings often fail to capture meaningful biology, displaying vector properties indistinguishable from those of randomly generated sequences. A key contributor to this performance issue is the models’ failure to learn the underlying biology from unevenly distributed sequence spaces in the training data.

Our novel, model-agnostic scoring framework is, to the best of our knowledge, the first to quantify protein sequence embedding reliability. We believe that our robust approach to screening embeddings prior to making biological inferences, stands to significantly enhance the reliability of downstream applications.

## Introduction

Language Models (LMs), originally developed for natural language processing (Qin et al., 2025), are increasingly accepted as the preferred *in silico* representation of protein, DNA, and RNA’s primary and higher-order structures (Consens et al., 2025; Wang et al., 2025; Weissenow & Rost, 2025; Zablocki et al., 2025). Their ability to learn an encoding that captures many aspects of a given biomolecule from simple amino or nucleic acid sequence, have made them a promising tool for deriving biological insights (Hoarfrost et al., 2022; Hugo et al., 2023; Lin et al., 2023; Marquet et al., 2022; R Prabakaran & Y Bromberg, 2025; Rives et al., 2021). LMs encode a biomolecule as an embedding, i.e. a sequence of numbers representing a point in a multidimensional latent space. Embeddings serve as powerful computational proxies for facilitating a range of downstream tasks, such as similarity searches, structural, functional annotations, and prediction of biomolecule properties (Dallago et al., 2021; Ji et al., 2021; Littmann et al., 2021; Thumuluri et al., 2022). For instance, embeddings from protein Language Models (pLMs) have been used to predict protein function, mutation effect, subcellular localization, etc., achieving performance that rivals or surpasses traditional methods (Bromberg et al., 2024; Dallago et al., 2021; Fenoy et al., 2022; Littmann et al., 2021; Tran et al., 2023). Additionally, fine-tuning pre-trained pLMs has been shown to enhance predictions across multiple additional biological tasks, underscoring the versatility of these models (Dickson & Mofrad, 2024; Schmirler et al., 2024; Weissenow & Rost, 2025).

Despite the advantages of embeddings as biomolecular representations, the reliability or confidence of an embedding remains largely unquestioned. Unlike most machine learning-based predictions that have a corresponding prediction probability/reliability score, a given embedding is not questioned as a representation of a protein any more than a protein sequence would be. Embeddings are low-dimensional representation of biomolecules in the LM’s latent space, with each vector element serving as a coordinate in the map of this space. Coordinates are learnt to encode the training data, while minimizing the loss associate with the training tasks (Ciernik et al., 2024). The model’s uncertainty or confidence in an embedding originates from the same sources as any of its predictions—the LM’s training process, optimized to reach a computationally feasible solution that balances task performance within cost and time constraints, rather than achieving complete learning or a globally optimal representation. In simple words, a model’s latent space is just one of many possible optimal mappings for the given training dataset and the training objective. Moreover, datasets may not comprehensively capture the full sequence space — a limitation that is, arguably, even more obvious for protein sequences than for human languages. As a result, each protein’s projection into the latent space carries an inherent uncertainty, raising the question of whether embeddings consistently place every protein in a biologically meaningful position, where functionally or evolutionarily related sequences remain nearby.

Protein embedding’s uncertainty propagates to downstream tasks. Consider a protein language model (pLM) that can generate embeddings to serve as representations of protein sequences to enable downstream tasks such as predicting protein subcellular localization, T_*s*_, or transferring functional annotations between proteins based on their embedding similarity, T_*f*_ (Dickson & Mofrad, 2024; Littmann et al., 2021; Schmirler et al., 2024). Most pLMs do not provide an explicit measure of model’s confidence or uncertainty, U(P_1_), associated with an embedding E_1_ for a given protein P_1_. Devoid of a score to indicate the biological information content in an embedding, it difficult to pre-screen embeddings or diagnose sources of error in downstream tasks like T_s_ and T_f_.

Traditional sequence alignment-based approaches offer a biologically interpretable foundation for inference placed on top of their captured information, leveraging the level of evolutionary conservation as a well-established measure of confidence. pLM embeddings, in contrast, lack the standardized framework for evaluating their biological relevance and robustness across the complete range of sequences, seen or yet unseen. This issue is particularly clear when embeddings are derived from heterogeneous sequence distributions, as model biases or incomplete training data may lead to representations that fail to encode meaningful biological information. Without a mechanism to recognize low-confidence embeddings, erroneous predictions propagate across applications misleading biological insights.

Establishing a systematic approach to assessing embedding quality, independent of model architecture, training paradigms, or downstream applications, would enable the pre-screening of embeddings, improve downstream task performance, and, more importantly, enhance the reliability and interpretability of pLM-based predictions. Additionally, such an approach would provide a means to identify underrepresented and poorly learned regions of the protein space, guiding further experimental and computational exploration. Outside of biology, i.e. in the general language model space, there have been a few efforts in this direction. For example, Tsitsulin *et al*. proposed four complementary metrics to capture different aspects of unsupervised learning-generated embedding quality (Tsitsulin et al., 2023). While no single metric was found to be universally optimal, coherence, i.e. a measure of alignment between embedding dimensions to canonical basis vectors, and stable rank, i.e. a soft rank proxy for the effective dimensionality of a set of representations, were shown to correlate strongly with downstream shallow model performance. Similarly, May *et al*. proposed the eigenspace overlap score as a selection criterion for evaluating the quality of compressed (lower-dimensional) word embeddings. However, assessment of the uncertainty of learned protein representations in a biologically meaningful way has not yet been performed.

In this work, we illustrate a robust model-agnostic, empirical approach to measure the uncertainty associated with protein embeddings and to assess the biological relevance of said embeddings. By proposing methods for pLM embedding evaluation, we seek to standardize, expand, and improve the usefulness of these models in moving biology forward.

## RESULTS AND DISCUSSION

In this study, we systematically evaluate the quality of pLM-derived embeddings using the Astral dataset, a curated collection of protein structures. Our analysis reveals that inconsistencies in the generated embeddings stem from biased learning processes, where non-uniform representation of sequence space in the training set leads to suboptimal encoding. To address this challenge, we propose a rigorous screening framework to assess the biological relevance of embeddings prior to their application in computational analyses. By establishing quality control measures, we aim to enhance the reliability of pLMs for biological inference, ensuring their robustness in real-world applications.

### Embeddings share the same defect as predictions

Few protein Language Model (pLM) designs provide a confidence score to assess the quality of their learning. One such model is Evolutionary Scale Modelling (ESM) 2 (Lin et al., 2023), which provides per-residue predicted Local Distance Difference Test (pLDDT) scores that describe how well the predicted structure is expected to align with a corresponding real protein structure (Lin et al., 2023); we thus chose ESM2 to illustrate embedding uncertainty. We collected embeddings, predicted structures, and corresponding pLDDT scores by running ESM2 on the Astral40 protein dataset, which consists of 15,117 unique SCOPe domains (**Methods & Table S1**) (Brenner et al., 1996; Brenner et al., 2000; Chandonia et al., 2004; Fox et al., 2014). We aligned (Zhang et al., 2022) ESM2 predicted structures with respective PDB structures (Brenner et al., 2000; Burley et al., 2019) to compute Template Modelling (TM) scores (Zhang et al., 2022; Zhang & Skolnick, 2005), a standard measure of goodness of structural alignment (**Methods**). We labeled the quality of predicted structures as very high, high, moderate and low-quality if the TM scores fell within [0.9,1], [0.7,0.9), [0.5, 0.7), and [0, 0.5) bins, respectively (**Figure 1A**). ESM2’s mean pLDDT scores for the predicted structures, correlated well with TM scores, as expected of different measures of structural similarity (Pearson ρ=0.84, Kendal τ=0.56).

**Figure 1.**
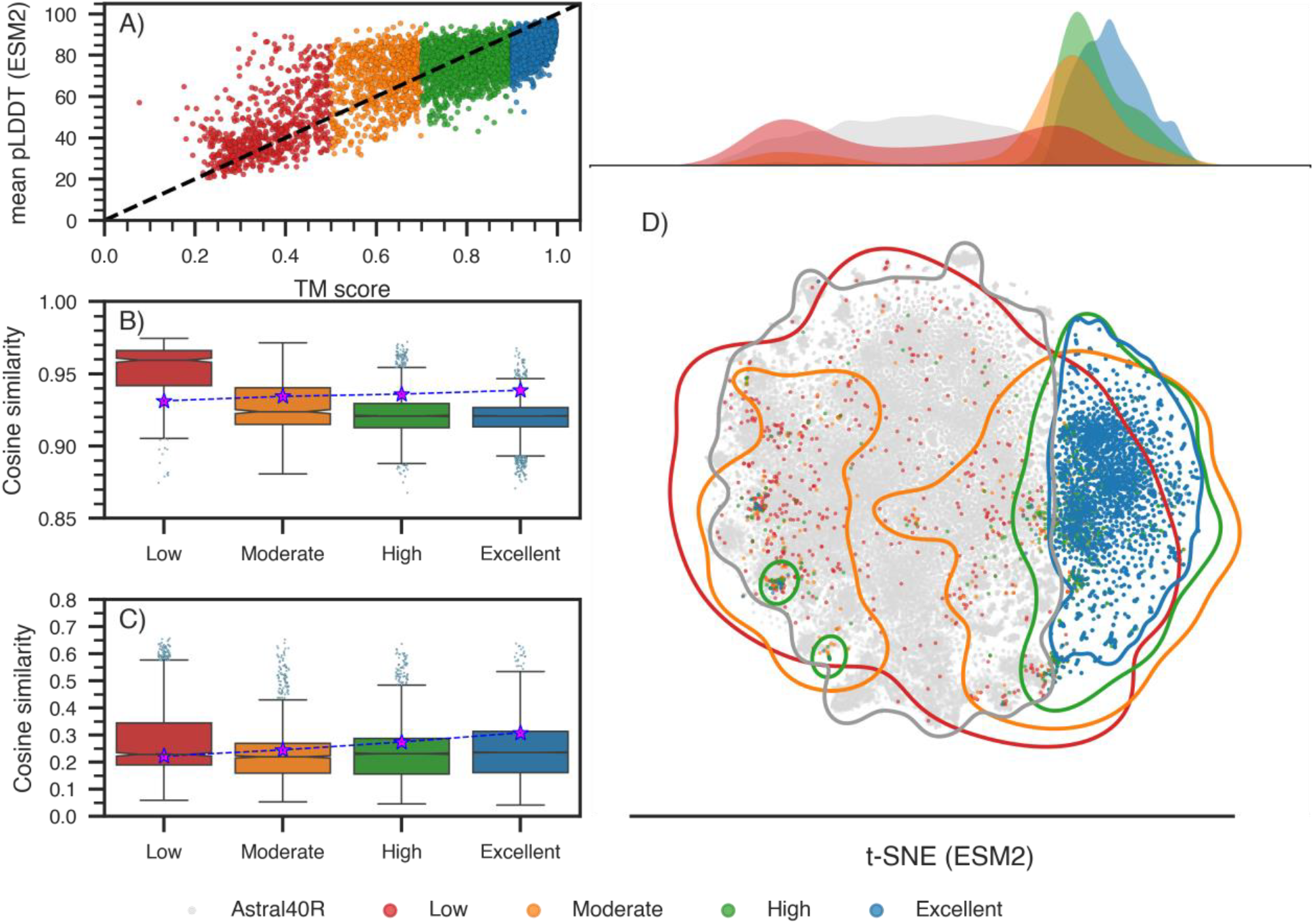
Structure prediction confidence and embedding distinctiveness. **(A)** ESM2 predicted structure quality (mean pLDDT, Y-axis) is, as expected, correlated with scores of predicted structure alignments with experimental structures (TM-score, X-axis) across Astral40 domains; here sorted by TM-score based confidence levels: Excellent (blue, TM > 0.9), High (green, 0.7 < TM ≤ 0.9), Moderate (orange, 0.5 < TM ≤ 0.7), and Low (red, TM ≤ 0.5). Cosine similarity between **(B)** ESM-2 and **(C)** ProtT5 Astral40 embeddings and a set of randomly generated, non-biological sequences (Astral40R). Purple stars denote the mean cosine similarity of embeddings within Astral40R, further highlighting the distinction between low and high-quality embeddings vs. random. **(D)** Two-dimensional t-SNE projections of ESM-2 embeddings for Astral40 (colored by TM-score category) and Astral40R (gray), illustrate the specifics of the overlap of the Astral40 embedding space with Astral40R. That is, Low scoring embeddings of real proteins (red lines) fall into a latent space also covered by random sequences (gray). At the same time, the latent space of Excellent scoring embeddings (blue) is nearly disjoint from the random space (gray)

We hypothesized that low-quality structures could derive from information-poor or ambiguous embeddings. To illustrate this, we created the *Astral40R* dataset that consists of five randomly shuffled sequences for every one of 15,117 unique domains in Astral40 (**Methods & Table S1**). Note, among the various methods for generating random sequence sets, we chose residue-shuffling because we hypothesize that one of the advantages of pLMs over traditional statistical models arises from their ability to learn more than merely the amino-acid composition of a sequence. We then computed the cosine and Euclidean similarities (**Eqn. 1**,**2**) of every domain/sequence in Astral40 (biological) against Astral40R (non-biological) datasets (**Figure 1B)**. Indeed, domain embeddings whose structures were accurately predicted were the least similar to embeddings of random sequences in Astral40R. As the quality of structures dropped, the similarity of domain embeddings to those of Astral40R increased – in other words, the embeddings corresponding to poorly predicted structures were less meaningful.

This observation is neither unexpected nor unique to the ESM2 model. We repeated our experiment with ProtT5 (Elnaggar et al., 2020) and other LMs (**Table 1**) and observed similar patterns (**Figure 1C and S1**), i.e. Astral40 sequences with low-quality ESM-predicted structures were not meaningfully represented by ProtT5 either.

**Table 1:** List of protein sequence embedders evaluated.

| Embedder | Model Size | Architecture | Embedding dimension | Trained dataset | Reference |
| --- | --- | --- | --- | --- | --- |
| ESM2 | 3B | Transformer | 2560 | UniRef50 | (Lin et al., 2023) |
| ProtT5 | 3B | T5 Transformer | 1024 | UniRef50 | (Elnaggar et al., 2020) |
| ProtTransT5BFD* | 3B | T5 Transformer | 1024 | BFD | (Elnaggar et al., 2020) |
| ESM1b* | 650M | Transformer | 1280 | UniRef50 | (Rives et al., 2021) |
| ESM1* | 650M | Transformer | 1280 | UniRef50 | (Rives et al., 2021) |
| Bepler & Berger* | 93M | BiLSTM | 121 | SCOPe | (Bepler & Berger, 2019) |
| PLUS-RNN* | 59M | RNN | 1024 | Pfam | (Min et al., 2021) |
| Word2Vec* | 6.4M | Skip-gram | 512 |  | (Asgari & Mofrad, 2015; Mikolov et al., 2013) |
| FastText* | 5.5M | Skip-gram | 512 |  | (Bojanowski et al., 2017) |
| Glove* | 6.4M | Matrix Factorization | 512 |  | (Pennington et al., 2014) |
| ESM2 (650)† | 650M | Transformer | 1280 | UniRef50 | (Lin et al., 2023) |
| ESM-1v† | 650M | Transformer | 1280 | UniRef50 | (Rives et al., 2021) |

\*Bio Embeddings was used to run these models (Dallago et al., 2021).
†These models were used only for the analysis of the Human variant effect prediction.

LMs are deep learning models, and the embeddings are as much model predictions as are the predicted structures. Predictions, however, are only as good as the model’s understanding of the input – in other words, if an input does not fall within the model’s scope, the prediction mirrors random inputs. We believe that this observation is common among all language models but, unlike ESM2, they are not trained to report a certainty (pLDDT-like) score that can be used to screen embeddings for future use.

### Is there a “junkyard” of embeddings?

We asked: is there a subspace for low-quality embeddings in pLMs’ latent spaces, where the low-quality, under-learned, noisy or less bio-meaningful embeddings go? We used t-SNE to explore model embeddings of Astral40 and Astral40R **(Methods)**. Indeed, we observed a distinct subspace for the random sequences of Astral40R in both ESM2 and ProtT5 latent spaces **(Figure 1D & S1A**).

We repeated this analysis for a select set of models covering diverse architectures (**Table 1**). We observed similar patterns for all ProtT5 family and ESM family pLMs, but not for Bepler and PLUS-RNN (**Figure S1 & S2**). The training tasks used for these latter two models differ from Mask Language Modelling (MLM), the standard training protocol for most LMs. PLUS-RNN is a Bidirectional recurrent neural network (RNN) model, trained with combination of MLM and contrastive learning loss, scoring similar representation of protein pairs from same Pfam families. Bepler is a bidirectional long short-term memory (LSTM) model trained on contrastive learning of global and local protein structural similarity. One could hypothesize the incorporation of contrastive loss sculptures latent landscape different from other LMs.

Traditional NLP algorithms word2vec, glove, and fasttext, which learn through the co-occurrence words or k-mers, failed to distance Astral40R embeddings from Astral40. Additionally, ESM2 embeddings of Astral40 biological sequences with high quality predicted structures (TM score ≥ 0.9; **Methods**) occupied a clearly distinct space within the Astral40 range, while sequence embeddings with low-quality structures spread out into the Astral40R random sequence space.

### What is a “good” embedding?

Based on these observations, we hypothesized that a protein positioned in the junkyard of a protein language model (pLM), i.e. in close proximity to non-biologically relevant sequences such as Astral40R, if the model fails to adequately learn its representation during training. As above, we define this region of poorly learned or uncharacterized protein embeddings as a “junkyard.”

We further propose that the degree of overlap in latent space between a protein’s nearest neighbors and the embeddings of non-biological sequences is inversely correlated with the model’s confidence in the embedding, i.e. the embedding quality. We formulate this relationship as **Random Neighbor score** (*RNS*_*k*_ (*P*_1_)), reflecting the number of non-biological sequence neighbors of a given protein *P*_1_in a given pLM’s latent space (**Eqn. 5)**. RNS is computed by measuring the fraction of randomly generated sequences among ‘k’ nearest neighbors of a given embedding (**Eqn. 5)**.

RNS is a measurable quantity reflecting the confidence of an embedding in a model’s latent space. We tested this new measure on the Astral40 dataset in ESM2’s latent space **(Figure 2A & B**). RNS correlated with the TM score of ESM2 predicted structures (vs. PDB structures) with a maximum Pearson correlation of -0.74 (Kendal τ =-0.41) across different values of k. For comparison, ESM’s pLDDT attained a correlation of 0.84 (τ =-0.56). Unlike the pLDDT score, however, which is an output of supervised learning, coupled with ESM2 architecture’s ability to predicted structures, RNS is model agnostic and (1) can be computed for any model immediately post model development, (2) does not depend on the downstream use of the embedding, and (3) could be further refined for the purposes of the data and tasks at hand. RNS depends on the parameter ‘k’, i.e. the number of nearest neighbors to consider. Higher k values are preferred due to the enhanced stability to perturbation and impact of outliers; however, k should be substantially smaller than the dataset size to ensure that the theoretical ideal RNS score =0. We observed a k in [200,1000] range to be appropriate for our study since the size of the datasets used in this study vary from 2,000 to 15,117 (**Figure 2B & Table S1**). Note, since RNS is computed as a fraction of random neighbors, we assume the biological, i.e. non-random, protein set to be diverse, as it acts as a secondary anchor for filtering out uncertain embeddings.

**Figure 2.**
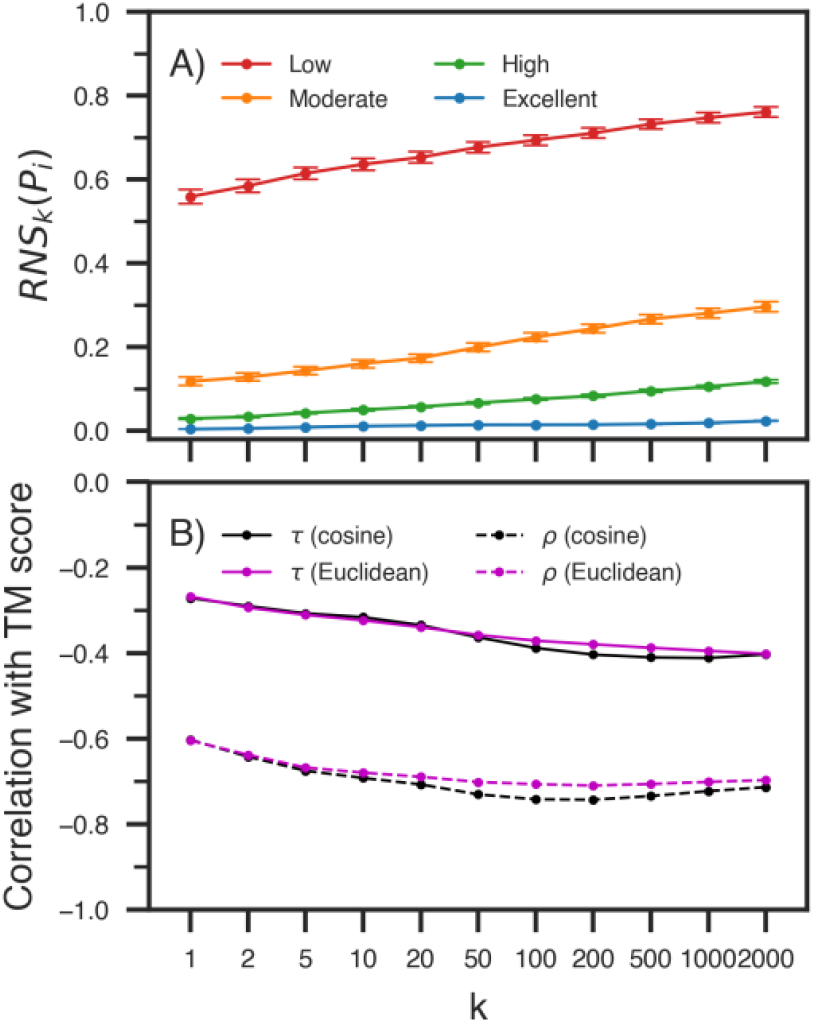
Random Neighbor Score (RNS) captures embedding uncertainty. **(A)** RNS (Y-axis), computed across varying values of *k* (K Nearest Neighbors, X-axis), effectively discriminates ESM-2 embeddings corresponding to low-confidence structures (TM ≤ 0.5, red line) from higher-quality ones (e.g. blue line – highest confidence structures). **(B)** RNS correlated with the TM-score, reaching a peak value (Pearson ρ=-0.74, Kendal t=-0.41) for higher ‘k’, reflecting its potential as a proxy for reliability in protein language model embeddings. RNS computed using Euclidean distance (magenta line) and cosine distance (black line) exhibit similar trends.

### Choosing a model for a protein set

Do learning biases make certain models more suitable for specific datasets? We applied RNS to evaluate the appropriateness of different pretrained protein language models (pLMs) for diverse protein sequence datasets. We compared the average RNS score (**Eqn. 6**) of different embedders (**Table 1**) on five datasets: **Astral40** domains, Intrinsically disordered proteins (IDP) and regions (IDR)(Aspromonte et al., 2024), novel metagenomic sequences (R. Prabakaran & Y. Bromberg, 2025), and novel “hallucinated” sequences (Anishchenko et al., 2021) (**Methods, Figure 3 & Table S1**). Note the term “Novel” in this context refers to sequences with <30% sequence identity to any protein in UniProt.

**Figure 3.**
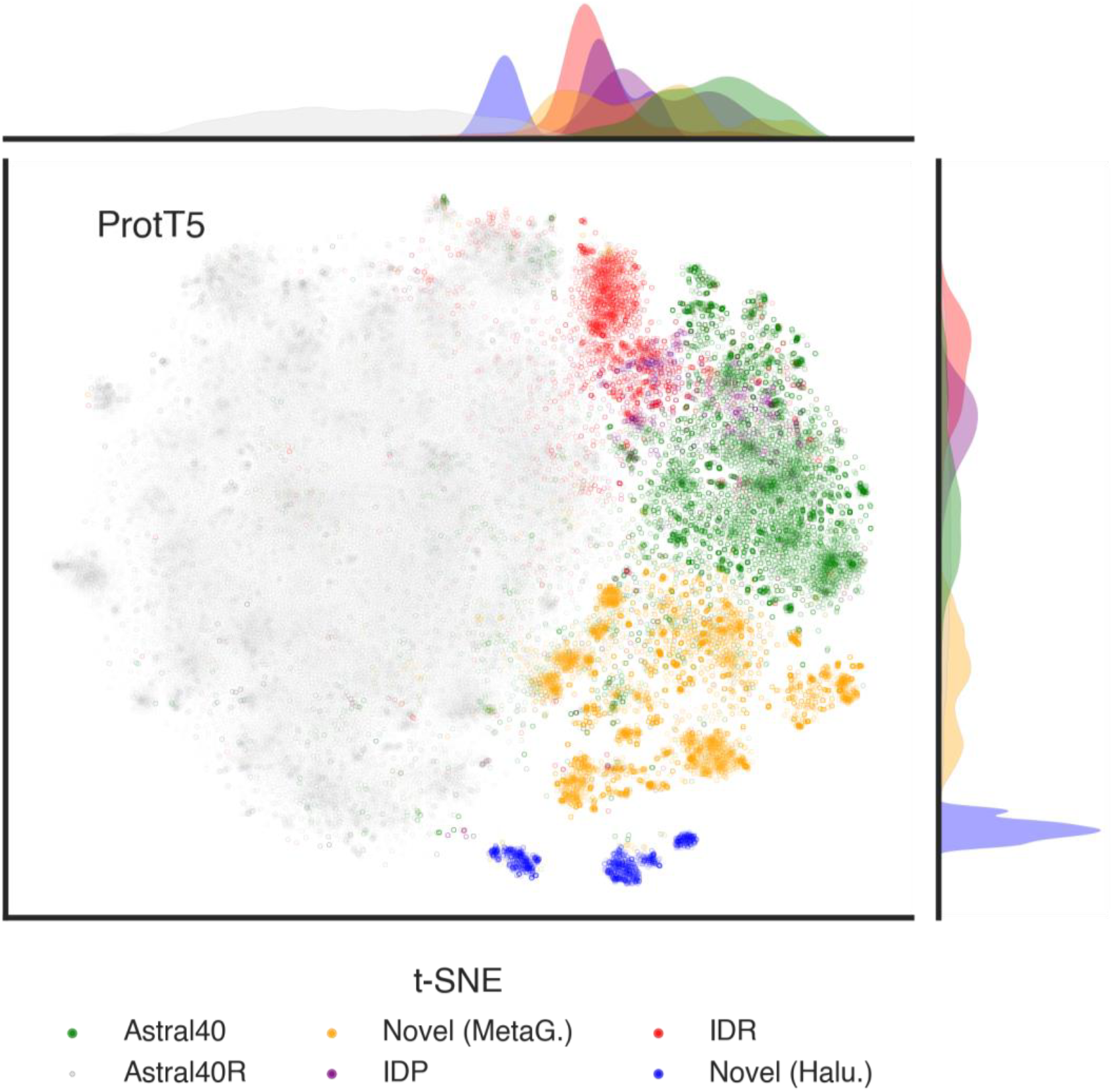
Uncertainty in pLM latent representations varies across protein sets. A two-dimensional t-SNE projection of the embeddings reveals distinct spatial distributions of diverse sequence sets, including: **(A)** protein domains from the Astral40 dataset, **(B)** Intrinsically Disordered Proteins (IDPs), **(C)** Intrinsically Disordered Regions (IDRs) of IDPs, **(D)** novel protein sequences derived from Metagenome-Assembled Genomes (MAGs), and **(E)** novel *de novo* Hallucinated Proteins (**Methods & Table S1**). (colors). The degree of overlap of each set with the non-biological/random Astral40R embeddings (gray) reflects the underlying uncertainty in the pLM’s (ProtT5, **Table S1**) latent space.

Among the tested models, ProtT5 and ESM2 consistently yielded the lowest RNS scores across all datasets (**Figure 4, Table S2**), suggesting higher confidence in their learned representations. However, ProtT5 (RNS_500=_0.12) significantly (*p-value* < 1E-6, Mann-Whitney U rank test) outperformed ESM2 (RNS_500_= 0.17) on Intrinsically Disordered Regions (IDRs, **Figure 4C**). This advantage may stem from ESM2’s training bias toward structured proteins, which would limit its ability to represent the disordered sequence space.

**Figure 4.**
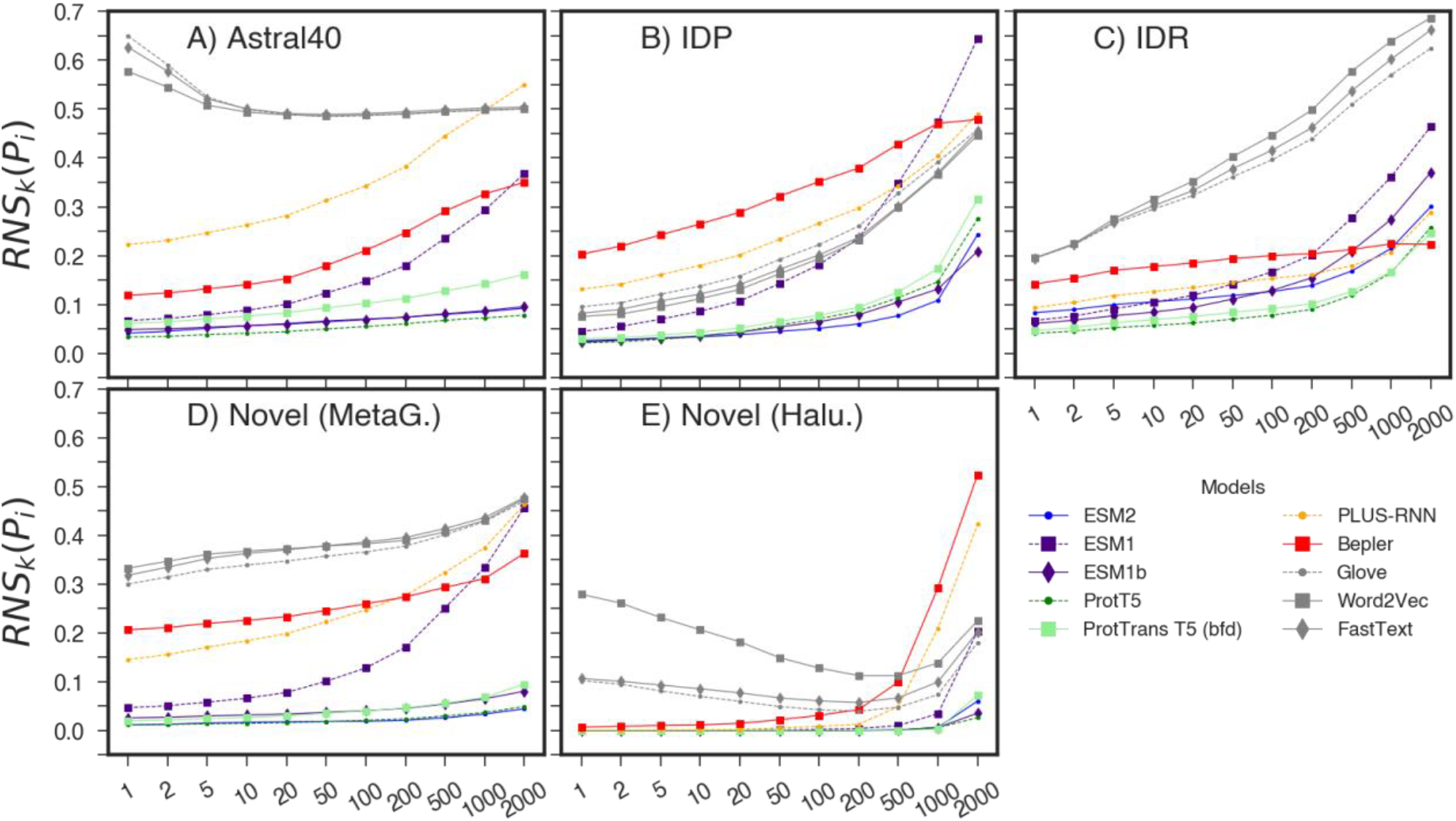
RNS measures embedding uncertainty across a range of protein sets and pLMs. RNS was computed for embeddings from multiple protein language models across diverse sequence sets, including: **(A)** protein domains from the Astral40 dataset, **(B)** Intrinsically Disordered Proteins (IDPs), **(C)** Intrinsically Disordered Regions (IDRs) of IDPs, **(D)** novel protein sequences derived from Metagenome-Assembled Genomes (MAGs), and **(E)** novel *de novo* Hallucinated Proteins (**Methods & Table S1**). Non-language model (Glove, Word2Vec and FastText) representations are colored in gray. RNS serves as a meaningful proxy for pLM’s uncertainty by measuring the degree of overlap with the non-biological/random Astral40R embeddings (Figure 3).

Interestingly, all models, including ProtT5 and ESM2, assigned higher RNS scores to IDRs compared to the structured Astral40 dataset, placing IDRs closer in representational uncertainty to their junkyards. Intrinsically Disordered Proteins (IDPs), which contain both structured and disordered regions, scored similarly to the Astral40 dataset, indicating partial model confidence.

Notably, NLP-based models, although generally underperforming in other datasets, showed improved performance on IDPs and IDRs. This may reflect the presence of low-complexity regions in IDRs, which produce distinctive k-mer signatures diverging from both Astral40 and Astral40R.

We also evaluated RNS on two novel sequence sets: metagenomic proteins, translated from predicted genes in metagenome-assembled genomes, and hallucinated sequences, generated *de novo* using TrRosetta. Surprisingly, most pLMs, except ESM1, PLUS-RNN, and Bepler, assigned low RNS scores to novel metagenomic sequences, suggesting recognition of coherent biological patterns despite their divergence from training data (**Figure 4D**).

An even more striking pattern was observed for hallucinated sequences, which were generated from the parameters of trRosetta neural network that was built to predict distances and orientations of residue pairs for a given protein family. We observed that these generated sequences were scored with low RNS by all models, including NLP-based ones (**Figure 4E**). This uniformity suggests a shared recognition of synthetic sequences that are biologically plausible.

### RNS captures vector properties of unlearned embeddings

To examine the information content of un(der)learned embeddings, we compared the information richness of the **Astral40** embeddings with those of their randomly shuffled counterparts (**Astral40R**). The informational richness of each embedding was quantified using four metrics (**Eqn. 7-11**): the (1) L2 norm of the full sequence embedding, i.e. the measure of magnitude or length of the embeddings; (2) covariance of residue embeddings across the full sequence (*COV*_*prot*_) and mean covariance among residues embeddings within all possible 15-residue fragments of the sequence 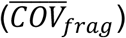, and (3) average cosine similarity among consecutive residues in all possible 15-residue fragments 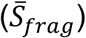. Residue covariance and Residue Similarity within a sequence indicate mutual agreement or disagreement between residues in vector space. Unlike randomly generated sequences, biological ones have evolved to perform specific functions. This constraint enforces synergy among residues for folding, dynamic conformational changes, allosteric regulation and carry out functions like catalysis.

We evaluated the difference in these metrics between embeddings of Astral40 with Astral40R datasets using standardized mean difference (**Cohen’s *d*, Eqn. 11, Figure 4A, Table S3**). While the L2 Norm significantly distinguished Astral40 from Astral40R for ESM family of models, this distinction was absent in other pLMs (**Table S3, Figure 5A**). In contrast, 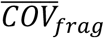 showed more consistent differentiation for the most evaluated pLMs, ESM family, PLUS-RNN and Bepler, but not the ProtT5 family. Interestingly, 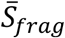 was significantly different between **Astral40** and **Astral40R** for ESM and ProtT5 families but not for PLUS-RNN and Bepler. These findings suggest that the different metrics capture distinct aspects of representational learning across architectures. To convey the signals of these diverse characteristics into a single uncertainty metric, we devised the Random Neighbor Score (RNS), which ranks each protein embedding’s position - relative to both randomized sequence and other protein embeddings - within the latent space of a pLM (**Figure 5B**).

**Figure 5.**
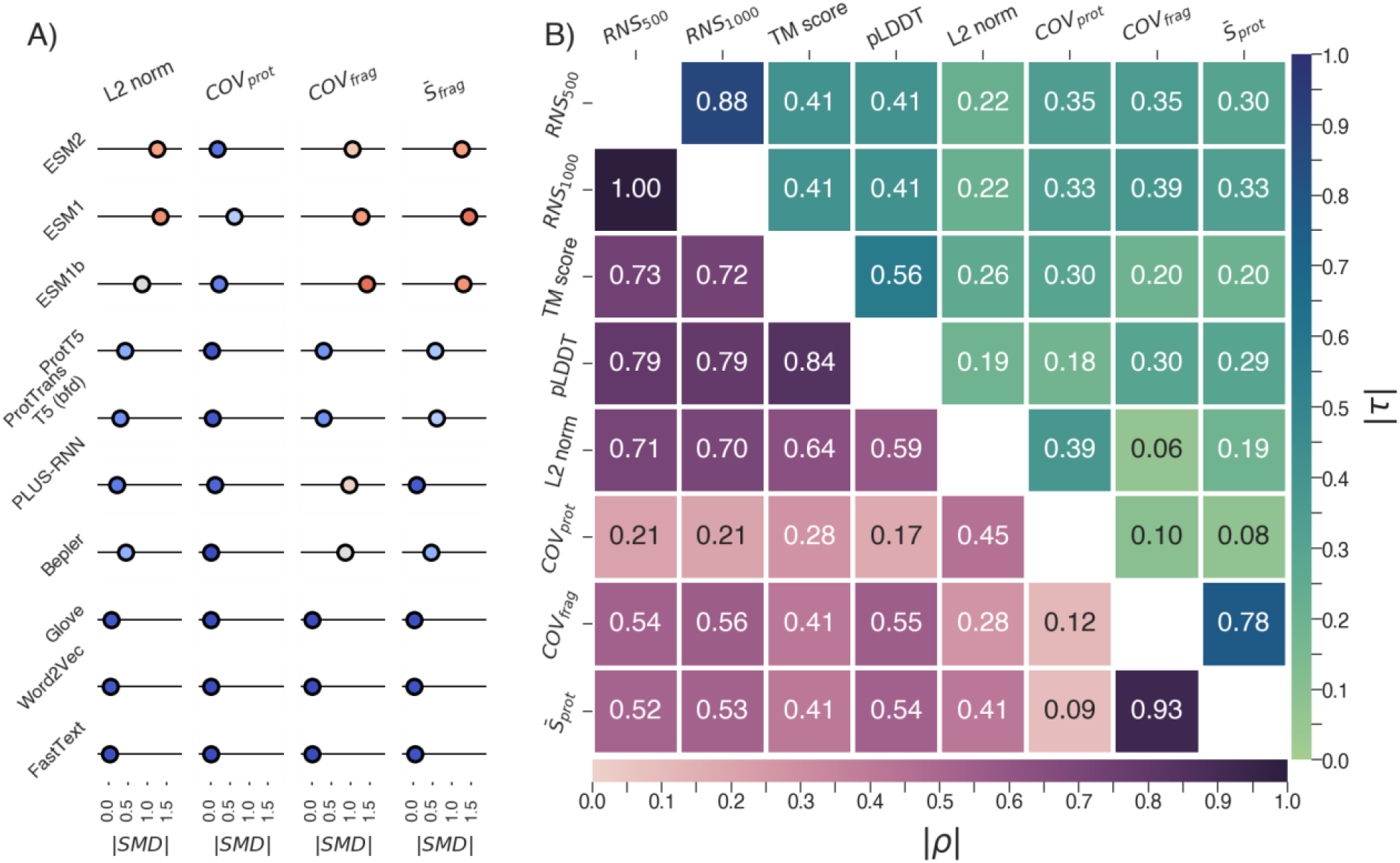
Information-theoretic measures as indicators of protein uncertainity. **(A)** The absolute Standardized mean differences (Cohen’s *d*, circle location on |SMD| X-axes) were calculated to quantify the distinction between embeddings of Astral40 domains and Astral40R random sequences across multiple information-content-based measures, revealing differences in sensitivities across protein language models (pLMs). The color of each circle ranges from blue (low SMD) to red (high SMD). **(B)** A Pearson correlation matrix presents the relationships between these measures and key confidence indicators, including Random Neighbor Score (RNS), ESM-2’s pLDDT scores, and the TM-scores comparing ESM2-predicted structures with experimental structures. Correlation coefficients are below the matrix diagonal, while Kendall rank correlations are displayed above the diagonal, highlighting both linear and rank-based associations.

We note that embedding representations of a protein sequence are derived from residue embeddings as either an average of all token (residue) embeddings or as a most representative token embedding. In either case, the complete set of token embeddings, an array of vectors, would be more informative. Indeed, in our analysis, *COV*_*prot*_ was less distinct than 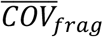, confirming the information loss incurred from averaging over the entire sequence. Nevertheless, protein-level embeddings remain widely used due to their convenience and lower computational costs. For this reason, we used protein-level embeddings to illustrate the utility of RNS in this study, while a more comprehensive residue-level analysis of RNS and related features may have been more informative.

### The un(der)learned portion of the human proteome informs predictive performance

We found that 19.1% (3,450) and 46.2% (8,372) of the human proteome (18,100 proteins of length ≤ 1,022, **Methods**) is un(der)learned (RNS_1000_ > 0) by ProtT5 and ESM2 (3B) models, respectively. Note that when all isoforms (n=37,477) are considered, these proportions increase to 19.7% (7,386) and 49.8% (18,677). Unexpectedly, ESM2 – a model with 3 billion parameters and also the backbone of ESMFold – exhibited substantially higher uncertainty than ProtT5. We hypothesize that this could stem from ESM2’s specialization for structure predictions, potentially at the expense of sequence-level generalization.

We extended our analysis to include other ESM variants (**Table 1)**, namely ESM1v and ESM2 (with 650M parameters). These demonstrated lower uncertainty of 27.5% and 15.2% of the human proteome, respectively. These results further underscore that larger models do not universally outperform smaller ones across tasks and datasets. Note that for this computation proteins exceeding 1,022 residues were excluded due to the sequence length limitations imposed by ESM models.

With a more relaxed RNS threshold (>0.1), illustrating the performance of a potentially more tolerant downstream application, the fraction of uncertain embeddings decreased, to 9.7% for ProtT5, 10.4% for ESM2 (3B), 8.8% for ESM2 (650M), and 16.1% for ESM1v, but did not disappear. Given that the human proteome is exceedingly well studied (Li & Buck, 2021), e.g there are many human protein structures in the PDB (Burley et al., 2019), the expectation for other proteomes would be a higher number of under-learned protein representations (Ding & Steinhardt, 2024). Thus, we expect that application of embedding screening prior to downstream applications could increase the precision of the latter.

To evaluate the above conjecture, here we demonstrate the utility of our RNS-based screening in improving the prediction of the effects of protein variants. We evaluated three pLMs: ProtT5, ESM1v, and the 650M-parameter version of ESM2 (**Table 1**) on two different datasets of human variants, as described in (Bromberg et al., 2024). Briefly, these are variants (Single Nucleotide Polymorphisms, SNPs) (1) annotated by PMD (Kawabata et al., 1999) as knock-out, effect, or neutral with regard to the corresponding protein function changes (*Function*) and (2) labeled by ClinVar (Landrum et al., 2014) as pathogenic or likely-pathogenic vs. identified as common or rare (Phan et al., 2020) in the human population (**Methods & Table S1**). For each model, we binned the proteins based on the RNS score of the corresponding protein and compared the loglikelihood score of variants, inferred from embeddings (Bromberg et al., 2024), to the variant classes. Note that although RNS can be computed at the residue level, this analysis focused on protein-level RNS for illustration purposes.

We evaluated the performance of these three pLMs in distinguishing SNPs as all combinations of [knock-out or effect] vs. neutral and [pathogenic or likely pathogenic] vs. [common or rare]. The models attained highest performance (AUROC>0.8, **Methods**) for variants from protein with proteins with zero embedding uncertainty (RNS=0; **Figure 6**); AUROC dropped to ∼0.5 for proteins with RNS>0.8.

**Figure 6:**
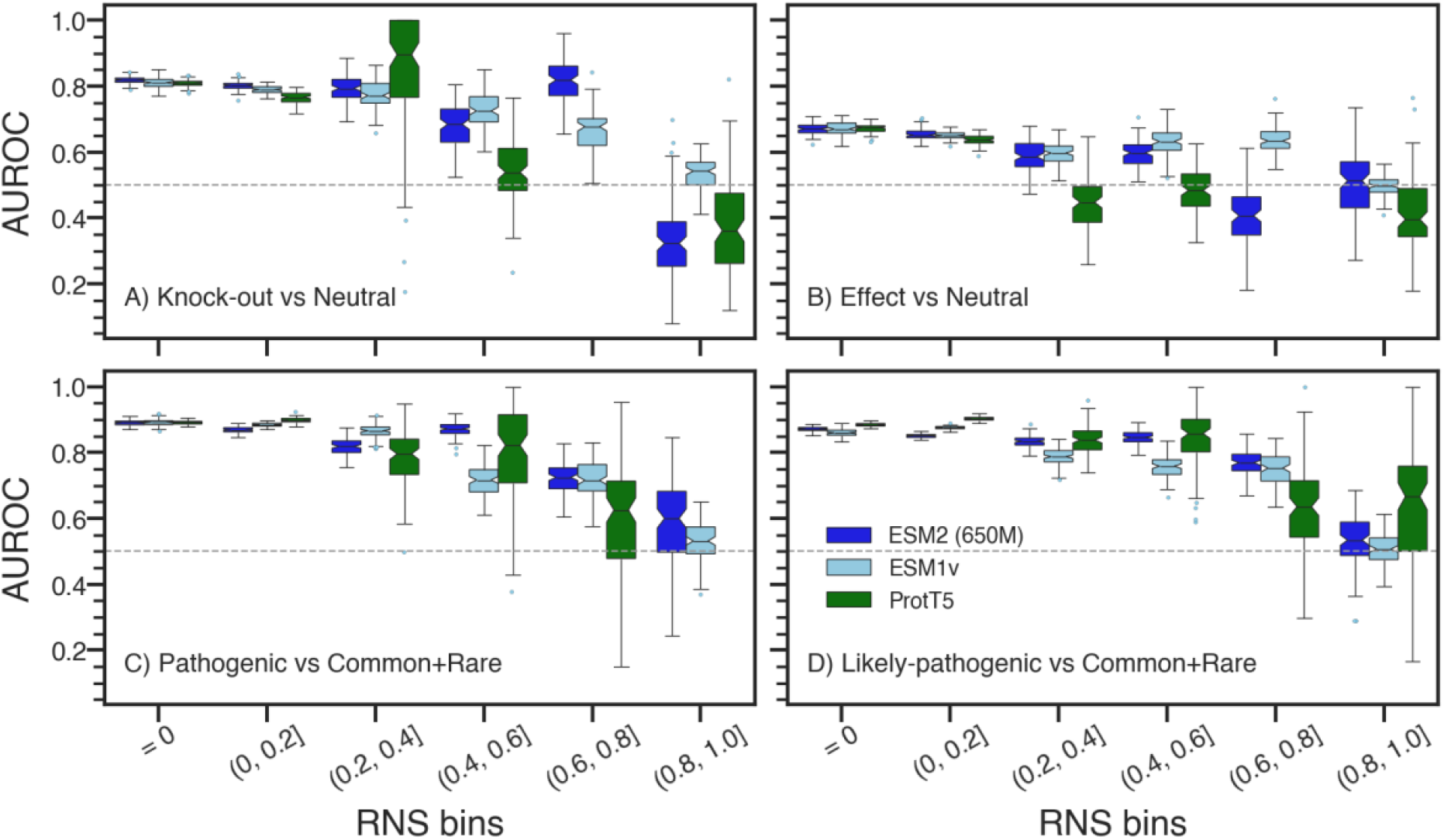
Variant classification tasks are improved with RNS-based screening. AUROC performance of three protein language models: ESM2 (650M parameters; dark blue), ESM1v (light blue), and ProtT5 (green), across binned Random Neighbor Scores (RNS) for four classification tasks: **(A)** Knock-out vs Neutral, **(B)** Effect vs Neutral, **(C)** Pathogenic vs Common+Rare, and **(D)** Likely-pathogenic vs Common+Rare. Each RNS bin (X-axis) left-to-right reflects increasing embedding uncertainty (higher values = greater uncertainty). Variant impact prediction performance declines at higher RNS, particularly beyond the 0.6 threshold, underscoring the relationship between embedding certainty and task performance. Horizontal dashed lines random classification (AUROC = 0.5). Blue circles represent statistical outliers beyond 1.5×IQR from the first and third quartiles of distribution.

Given that RNS quantifies the model’s uncertainty in its representation of a protein, we hypothesized an inverse relationship between RNS and the model’s ability to correctly identify functionally impactful variants. Across all three models, effects of variants in proteins with low RNS values (<0.5), i.e. more certain embeddings, were more indeed readily distinguishable.

Whereas, for proteins with higher RNS values, variant impact was indistinguishable from wild-type. We also observed this trend when comparing predictions of ClinVar pathogenicity vs. common SNPs (**Figure S4**).

These results demonstrate diminished predictive capacity in downstream tasks when using embeddings of greater representational uncertainty. In other words, pre-screening for embeddings of high uncertainty could improve model performance. Beyond improving existing model performance, we suggest that the ability to select well-represented samples could also lead to better training sets for newly-built downstream models. After all, better and more suitable-to-the-task data is arguably the best source of model improvement.

### All models are wrong, but some are useful

George Box’s famous maxim applies to protein language models (pLMs) just as well as any other model (George, 1976). Here we demonstrated the limitations of using pLM-produced protein embeddings as protein representations. We advocate for assigning confidence measures to these embeddings prior to any downstream applications. To this end, we propose several measures of embedding uncertainty estimates, reflecting incomplete learning or representation gaps. One of these, the Random Normalized Score (RNS) is a model-agnostic metric that can be readily applied to outputs of existing foundational models with minimal implementation effort. We argue that incorporating such screening strategies will enhance the reliability and interpretability of deep learning models in protein science. RNS can be integrated with pLM training workflows to pinpoint blind spots in the protein representation space and dynamically steer model training.

## MATERIALS AND METHODS

### Protein datasets

We collected 15,117 sequences and structures of SCOP domains of 20 to 1024 residues in length from the Astral40 dataset (Chandonia et al., 2004; Fox et al., 2014). Astral40 consists of non-redundant cluster representatives, clustered at 40% sequence identity. For every Astral40 sequence we generated five corresponding “synthetic” sequences by randomly rearranging the residues. Thus, we generated Astral40R, comprising 85,585 random sequences with same amino acid composition as Astral40 set.

In addition to Astral40 and Astral40R, we also collected four other sequence datasets: IDP (Intrinsically Disordered Proteins), IDR (Intrinsically Disordered Regions), Novel Metagenomic proteins and Novel Hallucinated proteins (**Table S1**). IDP and IDR consist of 2,030 and 4,146 intrinsically disordered proteins and regions, respectively, extracted from DisProt (Aspromonte et al., 2024). The Novel Metagenomic protein set consists of 11,444 proteins translated from genes of metagenome assembled genomes of <30% sequence identity to UniRef100 (Mitchell et al., 2020; R. Prabakaran & Y. Bromberg, 2025). The Novel Hallucinated protein set includes 2,000 synthetic proteins of length 100 residues, generated using the trRosetta deep neural network (Anishchenko et al., 2021).

### Variant dataset

We used two dataset of human variants: 1) the functional set consists of 3,818, 1,584, and 1,777 variants (Single Nucleotide Polymorphisms, SNPs) from 2,056 human proteins, classified as knock-out, effect, and neutral based on the effect of these variants on protein function (Bromberg et al., 2024; Kawabata et al., 1999) and 2) the pathogenic set comprising 2,499 and 4,804 pathogenic and likely-pathogenic variants from ClinVar, combined with 1,887 common (minor allele frequency, MAF ≥ 0.01) and 3,073 rare variants (0.01>MAF≥ 0.001) from 1,430 human proteins (Bromberg et al., 2024; Coordinators, 2013; Landrum et al., 2014). For each of the variants, we extracted the loglikelihood scores (Bromberg et al., 2024) from pLMs: ESM2 (650M), ESM1v and ProtT5 **(Table 1)**.

### Extracting embeddings from pLMs

We analyzed protein representations in the form of embeddings derived from a range of pretrained protein language models (pLMs). These included models from the ESM family—ESM-2 (3B and 650M), ESM-1b, ESM-1v, and ESM-1 (Lin et al., 2023; Rives et al., 2021)—and the ProtTransT5 family, comprising ProtT5 (ProtT5-XL-U50) and ProtTransT5BFD (Elnaggar et al., 2020). For comparison, we also included PLUS-RNN (Min et al., 2021), the Bepler & Berger model (Bepler & Berger, 2019), and three classical natural language models, Word2Vec (Mikolov et al., 2013), FastText (Bojanowski et al., 2017), and GloVe (Pennington et al., 2014), adapted for amino acid sequences.

Protein embeddings from all models, except for ESM-2 (3B & 650M), ESM-1v, and ProtT5, were generated using the BioEmbeddings framework (Dallago et al., 2021). Embeddings for ESM-2 and ESM-1v were obtained using the official esm Python library, while ProtT5 embeddings were retrieved via the Hugging Face Transformers interface. For all models and datasets, protein-level embeddings were computed by averaging residue-level embeddings across the full length of each sequence.

### Embedding similarity and distances

To quantify the similarity between two embeddings (e_1 &_ e_2_), both cosine and Euclidean similarity metrics were employed (**Eqn. 1, 2**). Similarly, we had used cosine and Euclidean distances to measure distances between embeddings (**Eqn. 3, 4**). Although all analyses were conducted using both distances/similarities, only the results derived from cosine-based measures are reported in this manuscript, given the consistency of the inferences obtained. Note that we also repeated all analyses using length normalized embeddings (unit vectors), observing results consistent with those obtained without normalization.

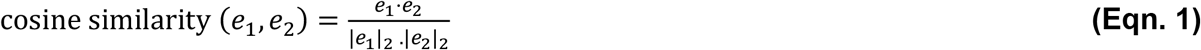

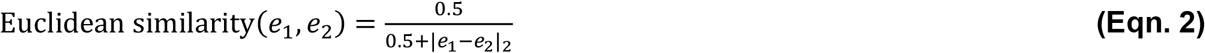

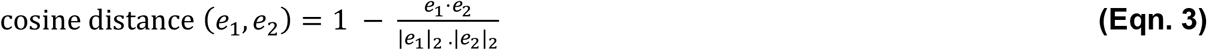

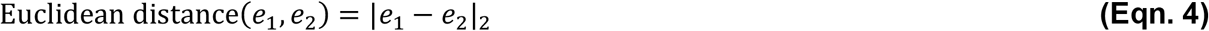

### Embedding visualization

We used t-SNE for dimensionality reduction of the pLM latent space. t-SNE 2D-projections were computed using (Policar et al., 2024) with perplexity of 45 for 150 iterations, followed by 200 iterations of exaggeration of 4, using cosine distance as metric.

### Random Neighbour Score (RNS)

We propose a new measure to quantify uncertainty of an embedding generated by a pLM, as a protein representation. Random Neighbor score of a protein P_1_ (*RNS*_*k*_(*P*_1_)) is the fraction of non-biological neighbors in a pLM’s latent space (**Eqn. 5)**. RNS is computed by measuring the fraction of generated sequences with random amino acid residue permutations among *K* nearest neighbors of a given embedding. RNS can further be employed to estimate fitness of a pLM for representation of a set of proteins (**Eqn. 6**).

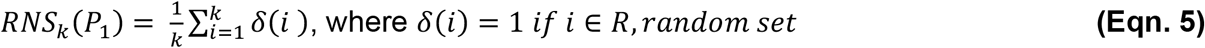

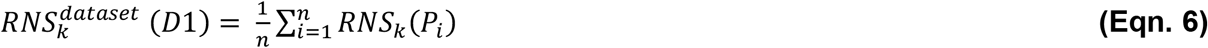

In this work, for a given set of proteins, the neighbors were identified from a combined pool comprising set protein embeddings and those of Astral40R, described above, i.e. a reference set of randomly generated non-biological sequences. To mitigate potential biases due to dataset imbalance, we applied under-sampling to ensure an equal number of embeddings from each sequence set in each of the 100 bootstrap iterations. The neighborhood size (*K*) was varied from 1 to 2000 to assess the sensitivity of RNS across scales.

### Features derived from Embeddings

To quantify the information content of protein embeddings, we computed four different measures:

1) L2 norm of the full sequence embedding, representing the embedding magnitude.

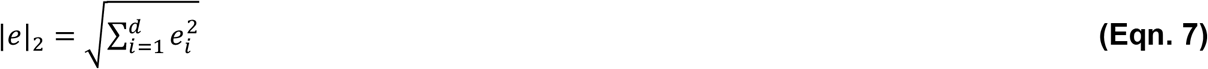
2) Covariance of residue embeddings across the full sequence (*COV*_*prot*_)

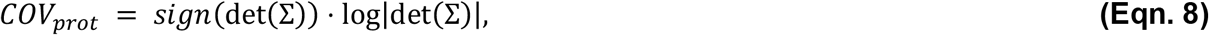

where 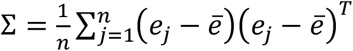 and n is the length of the protein and the number of residue embeddings.
3) Mean covariance among residue embeddings within all possible 15-residue fragments of the sequence 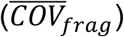

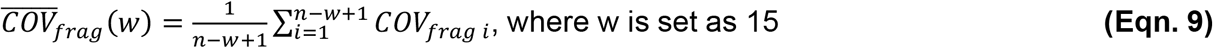
d) average cosine similarity among consecutive residues in all possible 15-residue fragments 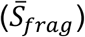. We introduce this Successive Residue Cosine Similarity (SRCS) to measure the change is a similarity between proximal residues along the protein sequence (**Eqn. 9**). SRCS is a vector of similarity values, described as

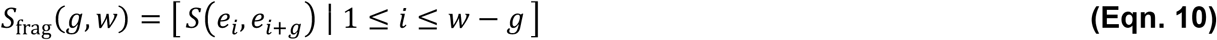

where S(e_i_, e_j_) represents the embedding similarity (**Eqn. 1, 2**), w is the fragment length, and g denotes the sequence distance between residues. Unless otherwise specified, default values for w and g are 15 and 1, respectively.

For a given protein, we consolidate SRCS as a single score by averaging over all fragments of length w (**Eqn. 10**).

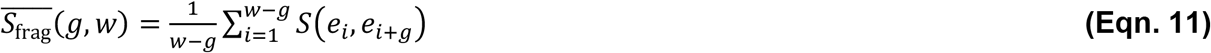

### Standardized Mean Difference

To quantify the size of the effect on embedding similarity of protein domains in real proteins vs. randomly generated non-biological sequences, we computed Standardized Mean Difference (SMD, Cohen’s *d*) for all embedding-derived metrics (**Eqn. 7-10**). For each metric (M), pLM, and Astral40 or Astral40R sequence origin. Cohen’s *d* was calculated as:

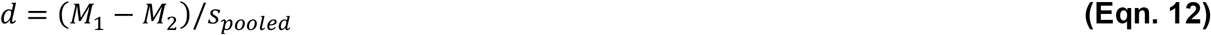

where M_1_ and M_2_ represent the mean values of the metric for the Astral40 and Astral40R sets, respectively, and pooled standard deviation (*s*_*pooled*_) is defined as:

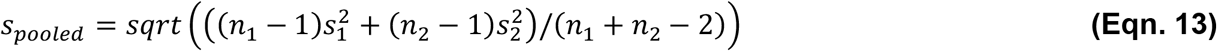

with s_1_ and s_2_ denoting the standard deviations, and n_1_ and n_2_ the sample sizes for each Astral40 and Astral40R.

Positive *d* values indicate that the metric is higher in Astral40 than in Astral40R, while negative values indicate the reverse. Effect sizes were interpreted using standard thresholds (small ≥ 0.2, medium ≥ 0.5, large ≥ 0.8); relative trends across pLMs were emphasized for comparative purposes.

### Evaluation on variant impact prediction

We measured the impact of RNS-based pre-screening of protein embeddings on variant impact prediction performance using standard metrics (**Eqn. 14-19**) and including Area under the Receiver Operating Characteristic curve (AUROC) and the Precision-Recall curve (AUPRC) (Fabian et al., 2011).

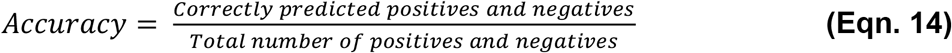

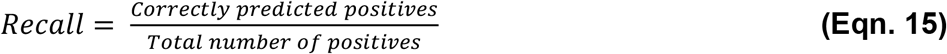

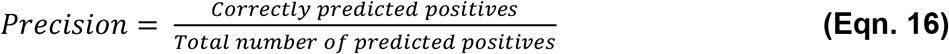

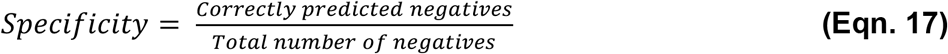

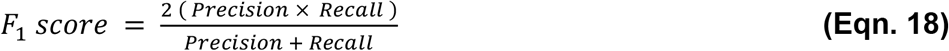

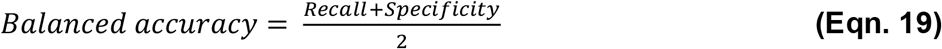

### Statistical tests

Statistical significances were assessed with the Mann-Whitney U rank test and Student’s t-test using SciPy (Virtanen et al., 2020).

## Supporting information

SOM

## Acknowledgements

This work was supported by the NSF (National Science Foundation) awards #2310114. We thank Dr. Ilya Nemenman for insightful discussions on the utility of protein embeddings, and Guintu Frederic D.M. for valuable input on the similarity of random embeddings.

## Coda and Data availability

A python package to compute RNS is available at https://bitbucket.org/bromberglab/rns/src/main/.

