## Supplementary material for "Quantifying uncertainty in Protein Representations Across Models and Task": SOM

**Prabakaran R[1*], Bromberg Y[1,2*]**

[1] Department of Biology, Emory University, Atlanta, GA 30322, USA
[2] Department of Computer Science, Emory University, Atlanta, GA 30322, USA

### Supplementary Information

#### Table S1: List of datasets used in this study

| Datasets | Type | Description | Number of sequences |
|---|---|---|---|
| Astral40 | Domains | Cluster representatives of the Astral dataset (SCOP domains), clustered at 40% sequence identity with length ranging from 20 to 1,024 residues (Fox et al., 2014) | 15,117 |
| Astral40R | Synthetic proteins (Random) | Five randomly generated (residues shuffled) sequences for every Astral40 sequence | 75,585 |
| IDP | Proteins | Proteins of length 20 to 1,000 residues containing disordered regions, extracted from DisProt (Aspromonte et al., 2024) | 2,030 |
| IDR | Fragments | Unique disordered regions, of length 24 to 997 residues, extracted from IDP set. | 4,146 |
| Novel metagenomic proteins | Proteins | Proteins of length 20 to 400 residues derived from metagenome assembled genomes from ESM Atlas (Lin et al., 2023). These proteins share <30% sequence identity to UniRef100 (Mitchell et al., 2020; Prabakaran & Bromberg, 2025). | 11,444 |
| Novel hallucinated proteins | Synthetic proteins | Synthetic 100-residue sequences generated by the trRosetta deep neural network, all of which are distinct from UniProt proteins (best BLAST E-value ≥ 0.1). (Anishchenko et al., 2021) | 2,000 |
| Functional | Protein variants | 3,818, 1,584 and 1,777 variants classified as knock-out, effect and neutral based on the effect of mutation on protein function (Bromberg et al., 2024) | 2,056 (7,179 variants) |
| Pathogenic | Protein variants | 2,499 & 4,804 pathogenic and likely-pathogenic variants, combined with 1,887 and 3,073 common and rare single variants (Bromberg et al., 2024) | 1,430 (12,263 variants) |

**Table S2: $RNS_{1000}$ values across datasets and pLMs**

| | Astral40 | Novel Hallucinated Proteins | IDP | IDR | Novel Metagenomic Proteins |
|---|---|---|---|---|---|
| **ESM2** | 0.113 | 0.055 | 0.155 | 0.345 | 0.045 |
| **ESM1** | 0.313 | 0.043 | 0.496 | 0.428 | 0.343 |
| **ESM1b** | 0.122 | 0.017 | 0.167 | 0.357 | 0.085 |
| **ProtT5** | 0.087 | 0.009 | 0.158 | 0.213 | 0.041 |
| **ProtTrans T5 (BFD)** | 0.176 | 0.013 | 0.220 | 0.254 | 0.083 |
| **PLUS-RNN** | 0.683 | 0.336 | 0.626 | 0.321 | 0.544 |
| **Bepler** | 0.657 | 0.762 | 0.907 | 0.505 | 0.684 |
| **Glove** | 0.973 | 0.533 | 0.891 | 0.654 | 0.745 |
| **Word2Vec** | 0.974 | 0.544 | 0.901 | 0.712 | 0.739 |
| **FastText** | 0.972 | 0.534 | 0.879 | 0.667 | 0.744 |

**Table S3: Standardized Mean Difference (Cohen's d) of Information-theoretic measures between structured domains and randomly generated sequences.**

| | L2 Norm | $COV_{prot}$ | $COV_{Frag}$ | $S_{frag}$ | $Dist_{Astral40}$ |
|---|---|---|---|---|---|
| **ESM2 (3B)** | 1.239 | 0.168 | 1.036 | 1.251 | 1.204 |
| **ESM1** | 1.321 | 0.619 | 1.265 | 1.427 | 1.165 |
| **ESM1b** | 0.841 | 0.218 | 1.429 | 1.296 | 1.360 |
| **ProtT5** | 0.405 | 0.034 | 0.294 | 0.553 | 1.444 |
| **ProtTrans T5 (BFD)** | 0.281 | 0.045 | 0.286 | 0.601 | 1.298 |
| **PLUS-RNN** | 0.193 | 0.101 | 0.953 | 0.066 | 0.840 |
| **Bepler** | 0.437 | 0.006 | 0.850 | 0.443 | 0.618 |
| **Glove** | 0.052 | 0.000 | 0.000 | 0.017 | 0.018 |
| **Word2Vec** | 0.043 | 0.000 | 0.000 | 0.023 | 0.028 |
| **FastText** | 0.018 | 0.000 | 0.000 | 0.032 | 0.026 |

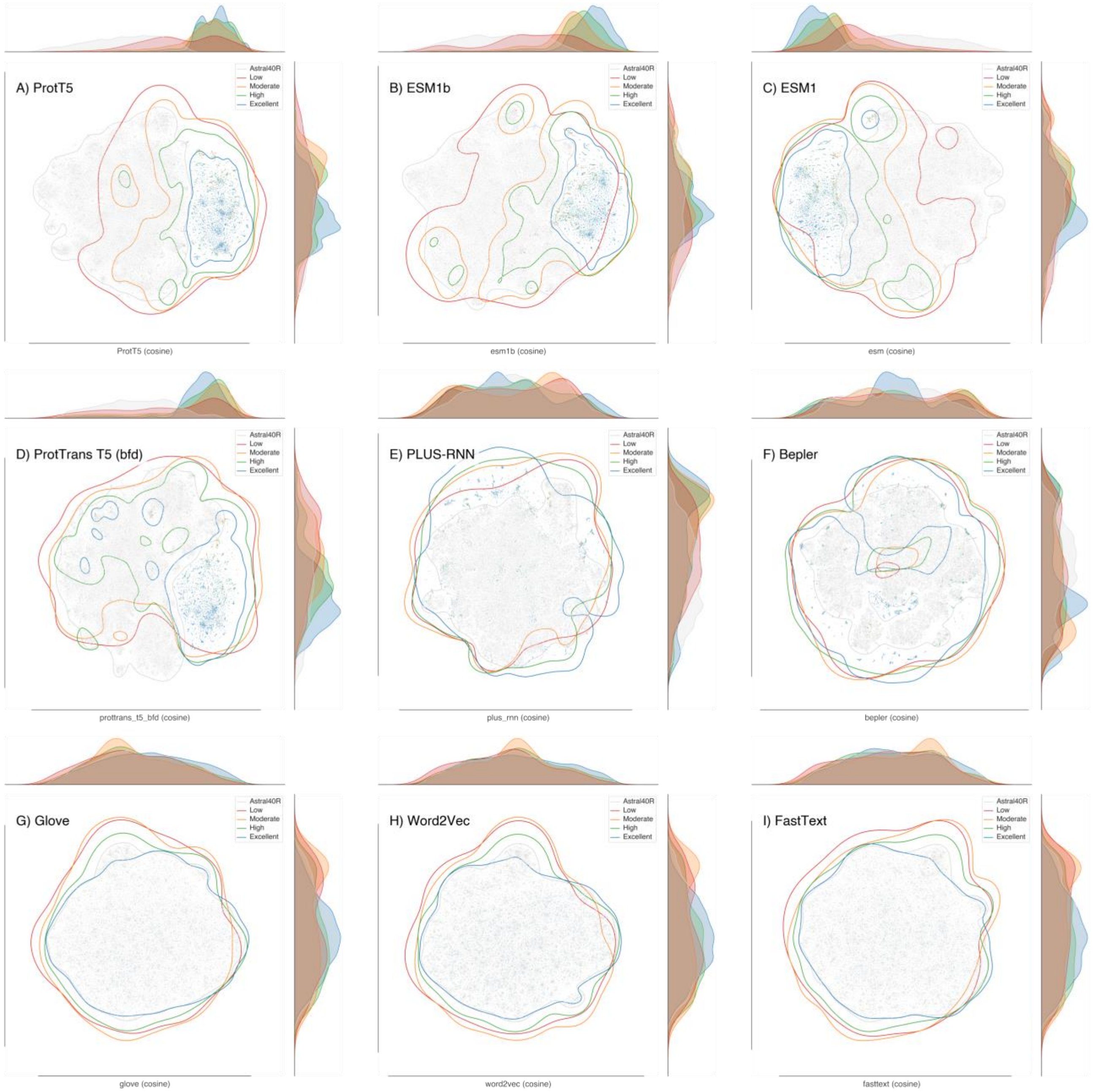


**Figure S1: Distinctiveness of biological sequences (Astral40 ) from randomly-generated sequences (Astral40R): t-SNE projection of latent space using cosine distance.** Latent space of pLMs listed in **Table 1** show boundaries between Astral40 and Astral40R embeddings. The Astral40 sequences are grouped (**Figure 1**) based on goodness of ESM's predicted structures (TM score).

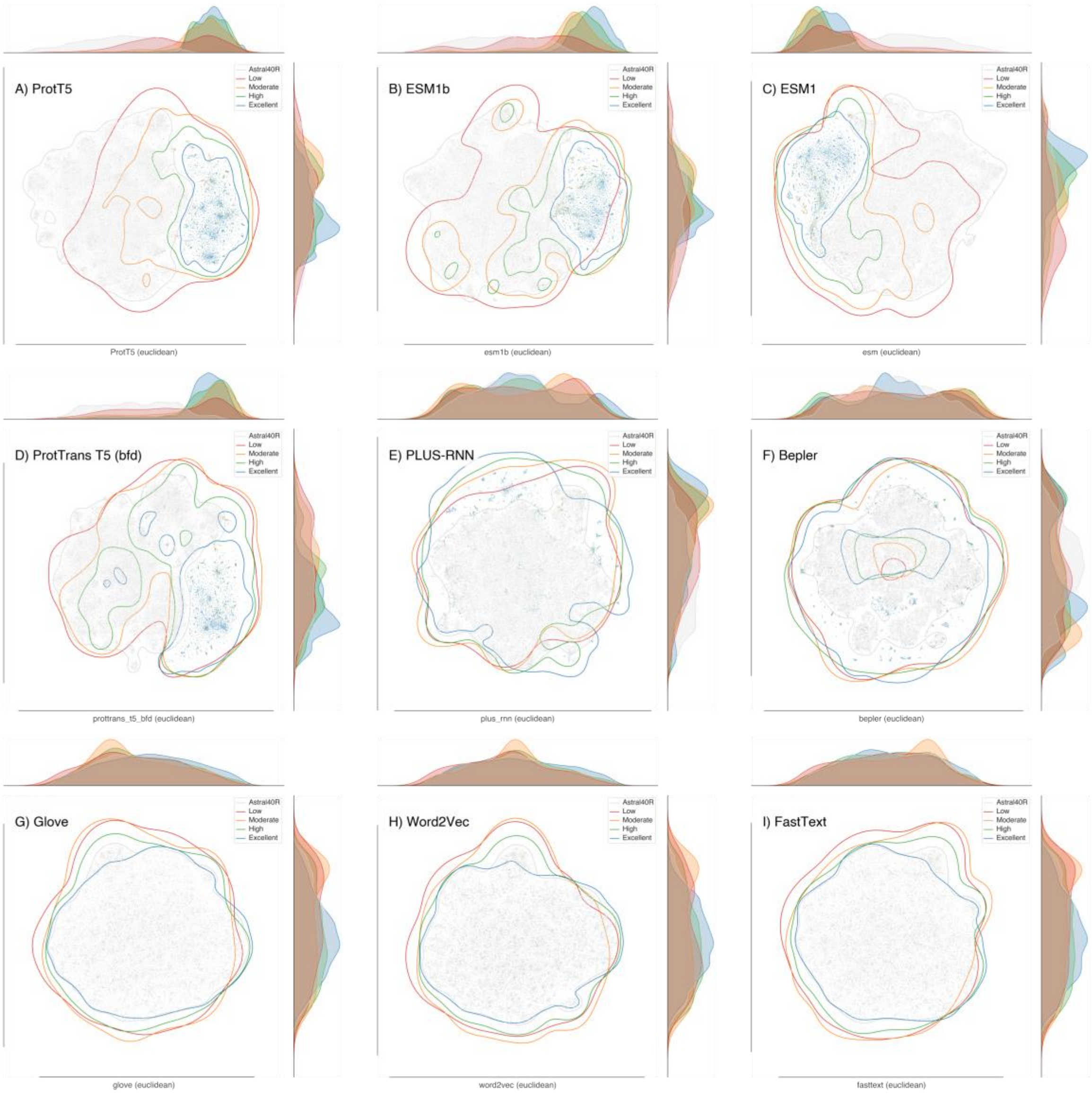


Figure S2: Distinctiveness of biological sequences (Astral40 ) **from** randomly-generated sequences (Astral40R): t-SNE projection of latent space using Euclidean distance. Latent space of pLMs listed in **Table 1** show boundaries between Astral40 and Astral40R embeddings. **The Astral40 sequences are grouped (**Figure 1**) based on goodness of ESM's predicted structures (TM score).**

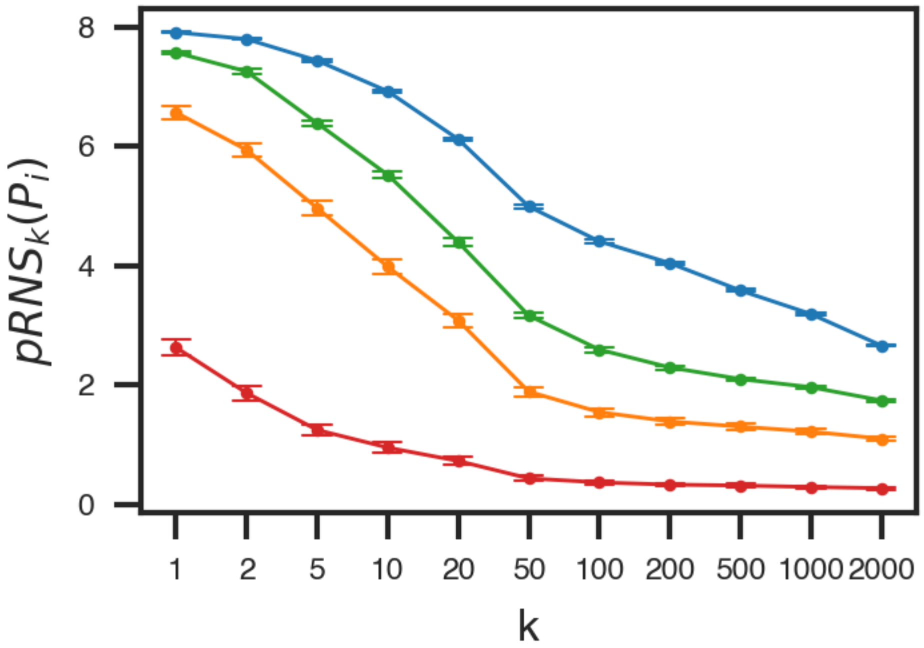


**Figure S3: Negative-Log transformed Random Neighbour Score for sensitive analysis**

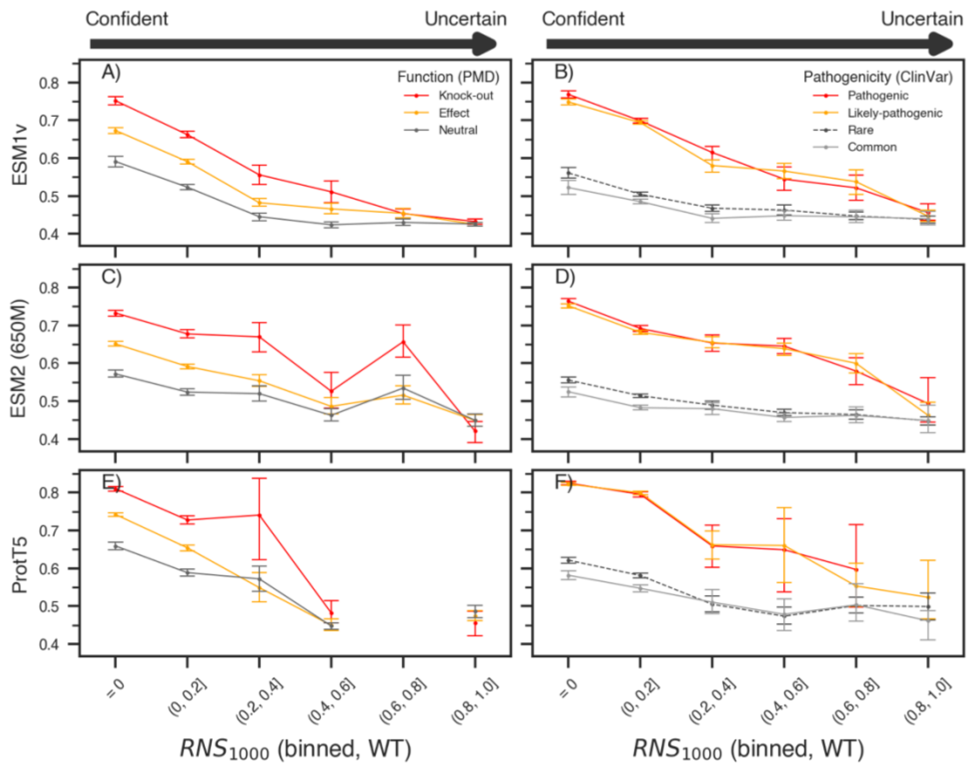


**Figure S4: RNS serves as the upper bound for the downstream task performance.** Unsupervised pLM-derived variant-effect scores for variants belonging to different functional (A, C & E) and pathogenic (B, D & F) variant classes are indistinguishable for protein with uncertain embeddings.
